# Deficiency of actin depolymerizing factors ADF/Cfl1 in microglia decreases motility and impairs memory

**DOI:** 10.1101/2024.09.27.615114

**Authors:** Sophie Crux, Marie Denise Roggan, Stefanie Poll, Felix C. Nebeling, Juliane Schiweck, Manuel Mittag, Fabrizio Musacchio, Julia Steffen, Katharina M. Wolff, Andrea Baral, Walter Witke, Christine Gurniak, Frank Bradke, Martin Fuhrmann

**Author notes:** **Corresponding authors:** Dr. Sophie Crux, Prof. Dr. Martin Fuhrmann, Neuroimmunology and Imaging Group, German Center for Neurodegenerative Diseases (DZNE), Venusberg-Campus 1/99, 53127 Bonn Germany. These authors contributed equally (Sophie Crux, Marie Denise Roggan, Stefanie Poll, Felix C. Nebeling).

## Abstract

Microglia are highly motile cells that play a crucial role in the central nervous system in health and disease. Here we show that actin depolymerizing factors ADF and Cofilin1 (Cfl1) are key factors of microglia integrity and function. We found a profound morphological phenotype in absence of ADF and Cfl1 in microglia. *In vivo* two-photon imaging of microglia with ADF/Cfl1-KO revealed reduced microglial fine processes motility and impaired microglia migration towards a laser-induced lesion. We found increased accumulation of stabilized F-actin and altered microtubule dynamics in ADF/Cfl1-KO microglia, indicating that ADF/Cfl1 are necessary for microglial cytoskeleton dynamics. Interestingly, microglial ADF/Cfl1-deficiency decreased learning and memory, suggesting that impaired microglial cytoskeleton dynamics affect neuronal functions relevant for cognition. Our results reveal a fundamental role of ADF/Cfl1 in microglia function and underscore the importance of these innate immune cells for higher cognitive functions.

## Main

Microglia act as resident immune cells of the central nervous system. They constantly scan the brain parenchyma in healthy brain and quickly react to sites of damage in an ATP-dependent manner^1,2^. These functions require an exceedingly dynamic rearrangement of the cell shape. The movement of very tiny microglial filopodia has been shown to be regulated by cAMP and depend on actin, but not microtubules^3^. The mechanisms, how the actin cytoskeleton regulates cell motility and migration are very well known^4,5^. However, how the function of the actin cytoskeleton relates to microglial motility and migration, especially *in vivo* in the brain, remains unknown.

In motile cells, actin filaments are constantly polymerized and depolymerized^5^. There exist three actin depolymerizing factor (ADF) isoforms that regulate actin dynamics and can sever actin filaments: ADF, cofilin 1 (Cfl1), and cofilin 2 (Cfl2). The three isoforms are present in many cells, but differently expressed throughout the body^6^. Cfl2 is the main isoform expressed in muscle^7^ and ADF/Cfl1 are the two isoforms that dominate functions in the brain^8^. In microglia, Cfl1 is highly expressed, ADF to a certain extend and Cfl2 is almost absent^9^. ADF and Cfl1 have been shown to be co-expressed in brain^8^ and suggested to have overlapping functions in brain development and physiology^8,10-12^. In the developing brain, Cfl1 is crucial for neuronal migration and cell cycle control, while ADF is dispensable during development^8^. Another important function of ADF/Cfl1 have been revealed for the regulation of axon growth in the adult spinal cord after injury^13^ and neurite growth in the developing brain^12^. Moreover, ADF/Cfl1 is implicated in synaptic physiology and vesicle trafficking^14^. In a mouse model of hemorrhagic brain injury, *Cfl1* siRNA knock down reduced microglia activation and improved the phenotype^15^. These findings underscore the importance of ADF/Cfl1 in the development and maintenance of the nervous system. However, research on the microglia-specific function of ADF/Cfl1 remains scarce.

Furthermore, microglia interact with different neuronal compartments by direct contact with their protrusions. It has been shown that neuronal activity influences microglia surveillance behavior^16^. In addition, microglia are crucial for synapse formation during learning and memory^17-19^. Studies investigating the role of microglial RhoA or Rac1, two proteins located upstream in signaling cascades of cofilin1, showed that in the absence of these proteins, microglia morphology and microglia-neuron interaction was impaired^20,21^. Interestingly, microglia with RhoA or Rac1 deficiency affected higher brain functions. To determine the role of ADF/Clf1 in microglia in relation to motility and migration, we used a conditional Cfl1-allele^7^ and combined it with the ADF knockout allele^8^ to generate a microglia-specific, ADF/Cfl1 double knockout line. Actin dynamics were crucially impaired in microglia of these mice, resulting in a striking morphological phenotype. Microglia did not move their fine processes anymore and were unable to migrate. In addition, this functional impairment of microglia had broader effects on cognition, since ADF/Cfl1 double knockout mice showed decreased learning and memory. These findings demonstrate the importance of an intact microglial actin cytoskeleton for their role as immune cells in relation to injury and cognition.

### ADF/Cfl1 deficiency results in aberrant morphology of microglia

In order to study the role of cytoskeletal proteins ADF/Cfl1 in microglia, we crossbred a quadruple transgenic mouse line (Cx3cr1ERT2^cre/wt^::Cfl1^fl/fl^::ADF^ko/ko^::Rosa26tdTomato^fl/wt^). This mouse line – abbreviated ADF/Cfl1-KO or KO – harbors a constitutive and ubiquitous *Adf* dominant negative mutation that leads to ADF deficiency **(Supplementary Fig. 1a-c)**. In addition, ADF/Cfl1-KO mice contain a conditional deletion of exon 2 in the *Cfl1* gene in Cx3cr1^+^ microglia **(Supplementary Fig. 1d, e)**, after tamoxifen-dependent Cre-recombinase activation. Lastly, Cx3cr1-specific tdTomato expression enabled microglia visualization via fluorescence. As control (WT), we chose the Cx3cr1ERT2^cre/wt^::Rosa26tdTomato^fl/wt^ mouse line **(Fig. 1a)**. To validate *Cfl1* knockout in microglia after tamoxifen application, we performed western blots of cultured primary microglia from ADF/Cfl1-KO mice. Tamoxifen treatment of primary microglia resulted in a downregulation of ADF/Cfl1 as validated by western blot and mRNA-expression analysis **(Supplementary Fig. 1f-h)**. Next, we performed a 3D morphological analysis of microglia in brain tissue comparing WT and ADF/Cfl1-KO mice. We intraperitoneally injected (i.p.) tamoxifen over 5 days and analyzed microglial morphology 11 days after the last tamoxifen application **(Fig. 1a)**. We prepared coronal brain slices containing the somatosensory cortex and performed fluorescent immunohistochemical stainings of microglia with an Iba1-antibody **(Fig. 1b, c)**. Comparing WT and ADF/Cfl1-KO mice, we found profound morphological alterations in microglia, such as significant soma enlargement and shortened microglial protrusions **(Fig.1d-f)**. 3D Sholl analysis of reconstructed microglia revealed a reduced cellular complexity, indicated by less intersections of branches with increasing distance in KO compared to WT microglia **(Fig.1 g, h)**. Thus, interfering with actin depolymerization dynamics in ADF/Cfl1-KO mice leads to a prominent morphological phenotype in microglia.

**Figure 1:**
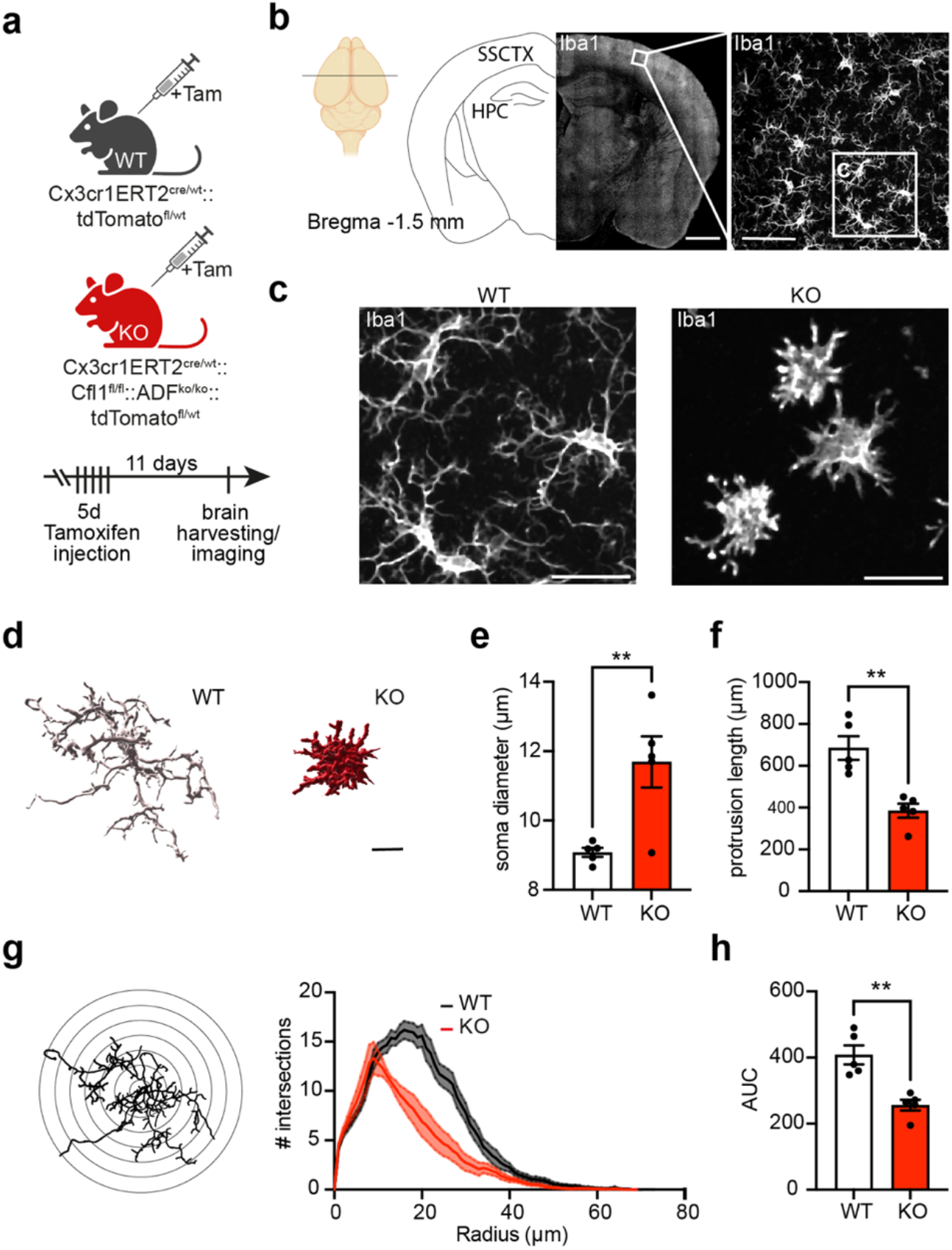
Morphological phenotype of microglia in absence of ADF/Cfl1. **a**: Transgenic mouse lines. WT: Tamoxifen inducible (ERT2) Cx3cr1-dependent tdTomato expression; KO: knock-out of ADF, inducible (ERT2) Cx3cr1-dependent KO of Cfl1 and tdTomato expression. Experimental paradigm with Tamoxifen injection. **b**: Schematic illustration of coronal brain sections for morphological analysis of microglia in the somatosensory cortex (SSCTX) above the hippocampus (HPC). Scale bars: overview of one hemisphere: 1 mm, zoom: 50 µm. **c**: Exemplary images of WT and ADF/Cfl-KO microglia. Scale bar: 20 µm. **d**: 3D reconstructed WT and KO microglia. Scale bar: 10 µm. **e**: Soma diaer of WT and KO microglia. Two-tailed t-test t=3.489, df=8, p=0.0082. WT n= 5; KO n=5 mice. **f**: Protrusion length per microglia comparing WT and KO. Unpaired two-tailed t-test, t=4.564, df=8, p=0.0018. WT n=5, KO n=5 mice. **g**: 3D Sholl analysis of WT and KO microglia. Exemplary reconstructed cell skeleton with radii for analysis demonstration. Distribution of # of intersections with processes in relation to radius from soma. **h**: Area under the curve (AUC) of Sholl curves of WT and KO, mean, unpaired two-tailed t-test, t=4.571, df=8, p=0.0018. WT, KO n = 5 mice/group.

### Motility of microglial processes is reduced in the absence of ADF/Cfl1

Microglia constantly scan the brain parenchyma by extending and retracting their processes, which likely requires a rapidly changing cytoskeleton. Hence, we investigated whether microglial process motility depended on actin remodeling regulated by ADF/Cfl1. We implanted a chronic cranial window on top of the somatosensory cortex, which allows *in vivo* two-photon (2P) imaging of tandem dimer Tomato (tdTomato) expressing microglia comparing WT and ADF/Cfl1-KO mice **(Fig. 2a)**. A three-dimensional volume (106 x 106 x 70 µm^3^) was imaged at a depth of 150 µm every 5 min over the time course of 25 min **(Fig. 2b)** to monitor dynamic changes of microglia protrusions (**Fig. 2 c, d**; **Supplementary Video 1**). Morphological changes – gained, lost and stable microglial protrusions – were measured at every 5 min time step and the 25 min average was calculated **(Fig. 2d, e)**. Microglia of ADF/Cfl1-KO mice had a smaller fraction of lost and gained protrusions in comparison to WT mice **(Fig. 2f, g)**. In contrast, stable microglial fractions were increased in ADF/Cfl1-KO compared to WT mice. **(Fig. 2h)**. Decreased loss and gain fractions in combination with a higher stable fraction resulted in a significantly reduced microglial process turn-over rate (TOR) in the absence of ADF/Cfl1 **(Fig. 2i)**. These data indicate a crucial role of actin depolymerization in microglial protrusion dynamics, the fundamental feature of microglial brain parenchyma surveillance.

**Figure 2:**
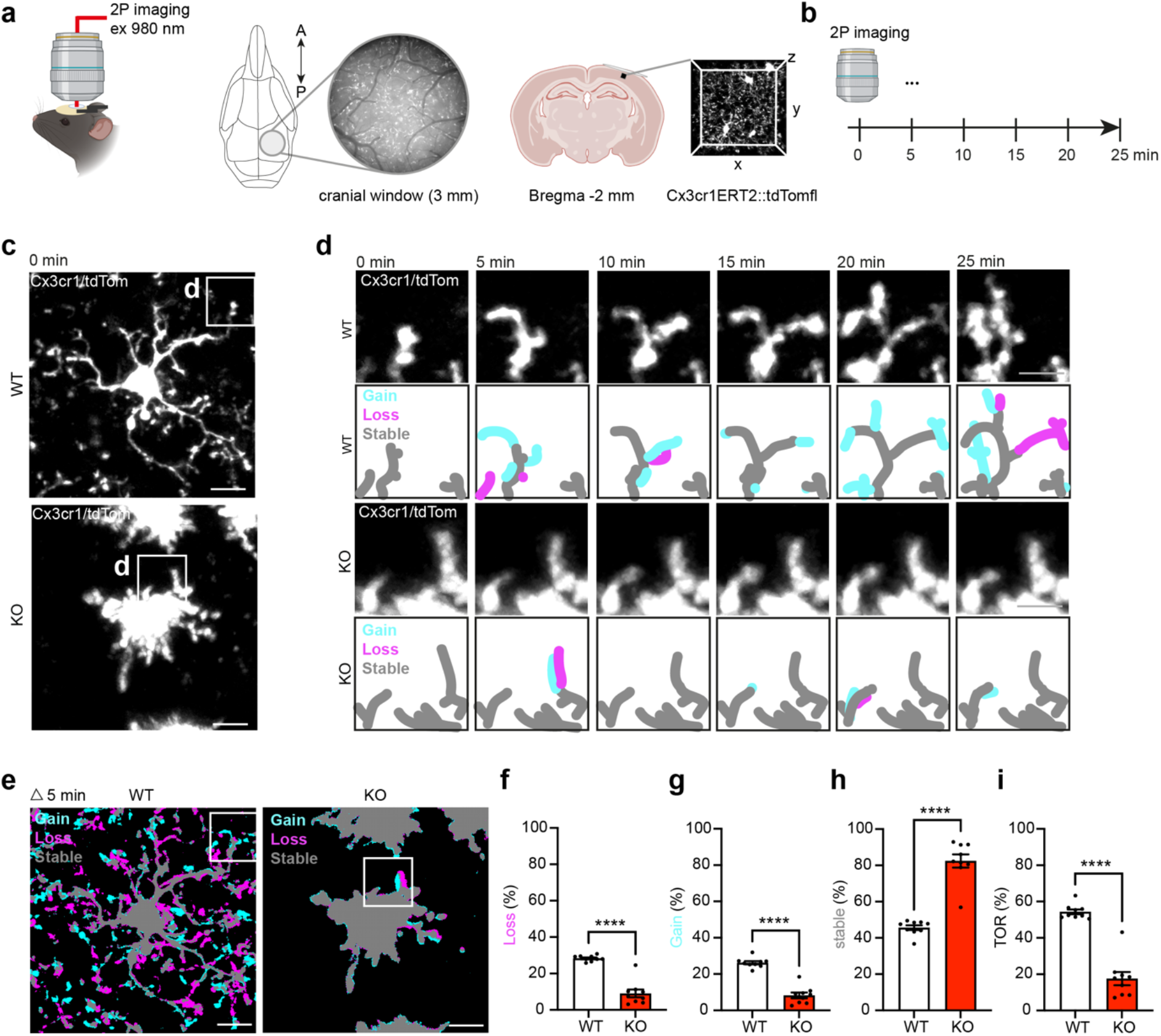
Impaired surveillance behavior of ADF/Cfl1-KO microglia. **a**: Experimental setup with chronic cortical window implantation and 2-photon imaging of microglia in a 3D volume in the somatosensory cortex. **b**: Imaging time scale over 25 min with 5 min interval. **c**: Z-projection of WT and KO microglia at 0 min. Scale bar 10 µm. **d**: Zoom from c with microglia protrusions and morphological changes every 5 min over 25 min. Protrusions and schematic drawing of changes over time of WT (top) and KO (bottom) microglia. Gain=cyan, loss=magenta, stable=grey. Scale bar 5 µm. **e**: Projection of WT and KO microglia with color-coded display of changes within 5 min. Scale bar 10 µm. **f-h**: Microglia signal in %, median of imaging time points. **f**: Fraction of loss. Two-tailed t-test, t=8.652, df=16, p<0.0001%. **g**: Fraction of gain, two-tailed t-test, t=9.808, df=16, p<0.0001. **h**: Fraction of stable. Two-tailed t-test, t=9.488, df=16, p<0.0001. **i**: Turn-over rate (TOR). Two-tailed t-test, t=9.488, df=16, p<0.0001. WT, KO n = 9 mice/group.

### Microglia migration towards brain injury requires ADF/Cfl1

Microglia, besides their constantly ongoing patrolling of the brain parenchyma, can rapidly migrate towards brain injuries^2,22-24^. To test whether this fundamental capability of microglia requires ADF/Cfl1, *in vivo* 2P imaging was performed in the somatosensory cortex through a chronic cranial window over 20 hours to monitor migrating microglia after laser lesion **(Fig. 3 a, b)**. In WT brains, microglia extended their protrusions until 8 h after lesion. 8–20 h after lesion they showed directed movement towards the site of injury **(Fig. 3 c, d; Supplementary Video 2)**. ADF/Cfl1 deficiency led to mostly stationary microglia with undirected to little movement and few sporadic extensions of protrusions towards the lesion site **(Fig. 3 e, f)**. Tracking of individual microglia within the time course of 8-20 h after lesion induction revealed that WT microglia started migrating at a radial distance of 140-250 µm and stopped within 140 µm from the lesion center **(Fig. 3 g)**. In contrast, ADF/Cfl1-KO microglia barely migrated or migrated in an undirected manner **(Fig. 3 g)**. The density of WT microglia increased within a 140 µm radius around the lesion center, whereas it decreased within the 140-250 µm segment. At a distance of more than 250 µm, microglia density did not change **(Fig. 3 h)**. In ADF/Cfl1-KO mice, the density of microglia decreased over time in all three segments, likely because of cell loss, since microglia did not migrate **(Fig. 3 h)**. We next calculated a directionality index (Di), which enabled us to quantitatively compare migration directionality between the two groups **(Fig.3i)**. WT microglia showed an almost ideal Di close to one (0.91 +/-0.01). In stark contrast, the DI of KO microglia was significantly smaller equal to zero (0.00 +/-0.06) **(Fig. 3j)**. These findings were corroborated by a small, but significantly reduced total travelled distance in conjunction with a strongly reduced Euclidian distance traveled when comparing WT and KO microglia **(Fig. 3k, l)**. These data suggested that the few microglia that migrated in ADF/Cfl1-KO mice travelled shorter distances and without any directionality towards the lesion. The reduced density of microglia in the three segments around the lesion center **(Fig. 3 g, h)** prompted us to systematically measure microglia density in our imaging volume prior to lesion induction. Indeed, we observed a reduced number of microglia in ADF/Cfl1-KO mice in comparison to WT **(Fig. 3 m, n)**. In summary, our data show that the physiological ability to extend protrusions and migrate towards a lesion site requires ADF and Cfl1 in microglia. In addition, ADF/Cfl1 deficiency severely affects microglia leading to cell loss.

**Figure 3:**
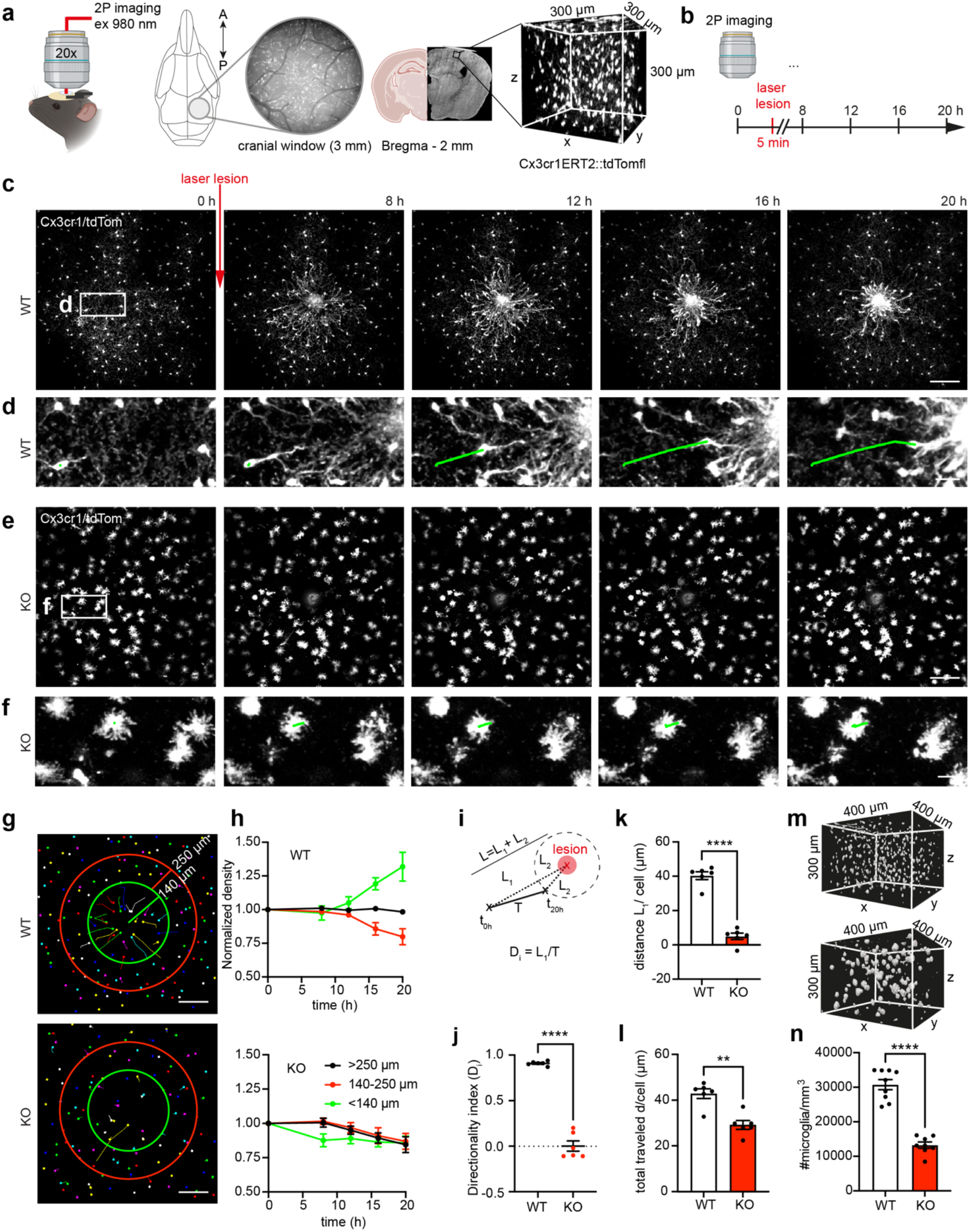
Impaired microglia migration in the absence of ADF/Cfl1. **a**: Experimental setup with chronic cortical window implantation and 2-photon imaging of microglia in a 3D volume in the somatosensory cortex. **b**: 3D imaging paradigm over 20 h including a laser lesion after the first volume (0 h). C/E: Exemplary projected images of microglia migration after laser lesion in WT (C) and KO (E) over 20 h. Scale bar: 100 µm. **d/f**: Close up of one tracked WT microglia extracting processes and migrating towards the lesion (**d**) and one tracked KO microglia not showing directed movement (**f**). Green line: tracked travelled distance. Scale bar: 20 µm. **g**: Exemplary image of tracked microglia 0 h – 20 h. Each dot (and line) represents one microglia. Red (250 µm) and green circles (140 µm) depict diameters around the lesion site. Scale bar 100 µm. **h**: Mean of normalized microglia density changes within diameters (140, 250, >250 µm) in 0 h - 20 h with SEM in WT (upper graph, n=6) and KO (lower graph, n=6). **i**: Schematic illustration of calculations of directionality index (Di). **j**: Di comparing WT and KO mice; unpaired two-tailed t-test, t=15.72, df=10, p<0.0001. WT, KO n = 6 mice/group. **k**: Average distance travelled (L1) of individual microglia into direction of the lesion. Unpaired two-tailed t-test, t=12.1, df=10, p<0.0001. WT, KO n= 6 mice/group. **l**: Average of total distance (d) travelled of individual microglia between t0 – 20 h. Mann-Whitney test U=1, sum of ranks 56,22; p=0.0043. WT, KO n = 6 mice/group. **m**: Exemplary *in vivo* 2P-imaged 3D volumes of rendered WT and KO microglia. **n**: Average microglia densities WT vs KO measured in 3D volumes. Unpaired two-tailed t-test, t=9.856, df=15, p<0.0001. WT, KO n = 9, KO n = 8 mice.

### ADF/Cfl1 shape the cytoskeleton beyond actin structures

We showed that ADF/Cfl1 as actin depolymerizing factors are crucial for remodeling and reorganization of the microglia shape, process motility and migration. Next, we further investigated the cytoskeleton in the absence of ADF/Cfl1 by directly staining filamentous actin (F-actin) with Phalloidin in fixed brain slices **(Fig. 4a, b)**. We observed a punctate staining within tdTomato positive WT microglia. On the contrary, in ADF/Cfl1-KO mice, F-actin strongly accumulated inside microglia, affecting more than 80% of microglia **(Fig. 4b, c)**. In complementary experiments using primary microglia, we confirmed that ADF/Cfl1 deficiency led to higher F-actin levels **(Fig. 4d-f)**. As in in *in vivo* conditions, primary microglia lacking ADF/Cfl1 changed their morphology, as most of the cells displayed an apolar shape. In contrast, vehicle-treated microglia had a normal polarized morphology **(Fig. 4g)**. To further characterize the distinct morphology of microglia in absence of ADF/Cfl1, we analyzed microglia expressing tdTomato using the FracLac plugin for Fiji. This plugin allows to describe cellular complexity regarding a variety of parameters, including circularity of the cell, span ratio, density and fractal dimensions. ADF/Cfl1-deficient in comparison to vehicle-treated microglia were more circular, had an increased density, a reduced span ratio and were less complex as indicated by the fractal dimensions **(Supplementary Fig. 2)**. To investigate changes of individual actin filaments on the sub-cellular level, we performed Airyscan super resolution microscopy **(Fig. 4h)**. While vehicle-treated microglia displayed contained F-actin puncta and few long, stable protruding filaments **(Fig. 4i)**, ADF/Cfl1-deficient microglia displayed actin arcs and stable fibers close to their cell border **(Fig. 4j)**. Live cell imaging of F-actin with SiR-actin confirmed the presence of increased stable actin filaments in microglia lacking ADF/Cfl1. In contrast, vehicle-treated cells predominantly contained F-actin puncta with few long actin filaments **(Supplementary Movie 3)**. Different cytoskeletal components interact with each other directly and indirectly to enable a plethora of cellular functions, such as motility of cells and establishment of cell shape^25^. Given the altered morphology of ADF/Cfl1-deficient microglia and the presence of actin arcs along the cell border, we asked whether other cytoskeleton components, such as microtubules, are affected by aberrant F-actin accumulation after ADF/Cfl1 deletion. Indeed, live cell imaging of SiR-Tubulin-labeled microglia revealed that microtubule protrusions were clearly reduced in microglia lacking ADF/Cfl1 **(Fig. 4k-m, Supplementary Movie 4)**. While ADF/Cfl1-deficient microglia form membrane protrusions in different directions, it appears that microtubules fail to invade into these protrusions, which is followed by a rapid membrane collapse, generating a spinning movement around the cell’s own axis **(Supplementary Movie 5)**. We further investigated this spinning movement and performed live cell imaging of tdTomato-expressing microglia for 3 hours. Indeed, microglia lacking ADF/Cfl1 moved less distance and rotated more frequently around their own axis **(Supplementary Fig. 2, Supplementary Movie 6)**. In conclusion, if the two actin depolymerizing factors ADF/Cfl1 are missing, F-actin strongly accumulates within microglia. Complementary to our *in vivo* findings, primary microglia in culture recapitulate changes in morphology and movement. On the sub-cellular level, we show that the morphological changes are accompanied by an increase in stable actin filaments and formation of actin arcs at the cell border. Additionally, decreased protrusion of microtubules and undirected cell movement point towards a widely affected cytoskeleton.

**Figure 4:**
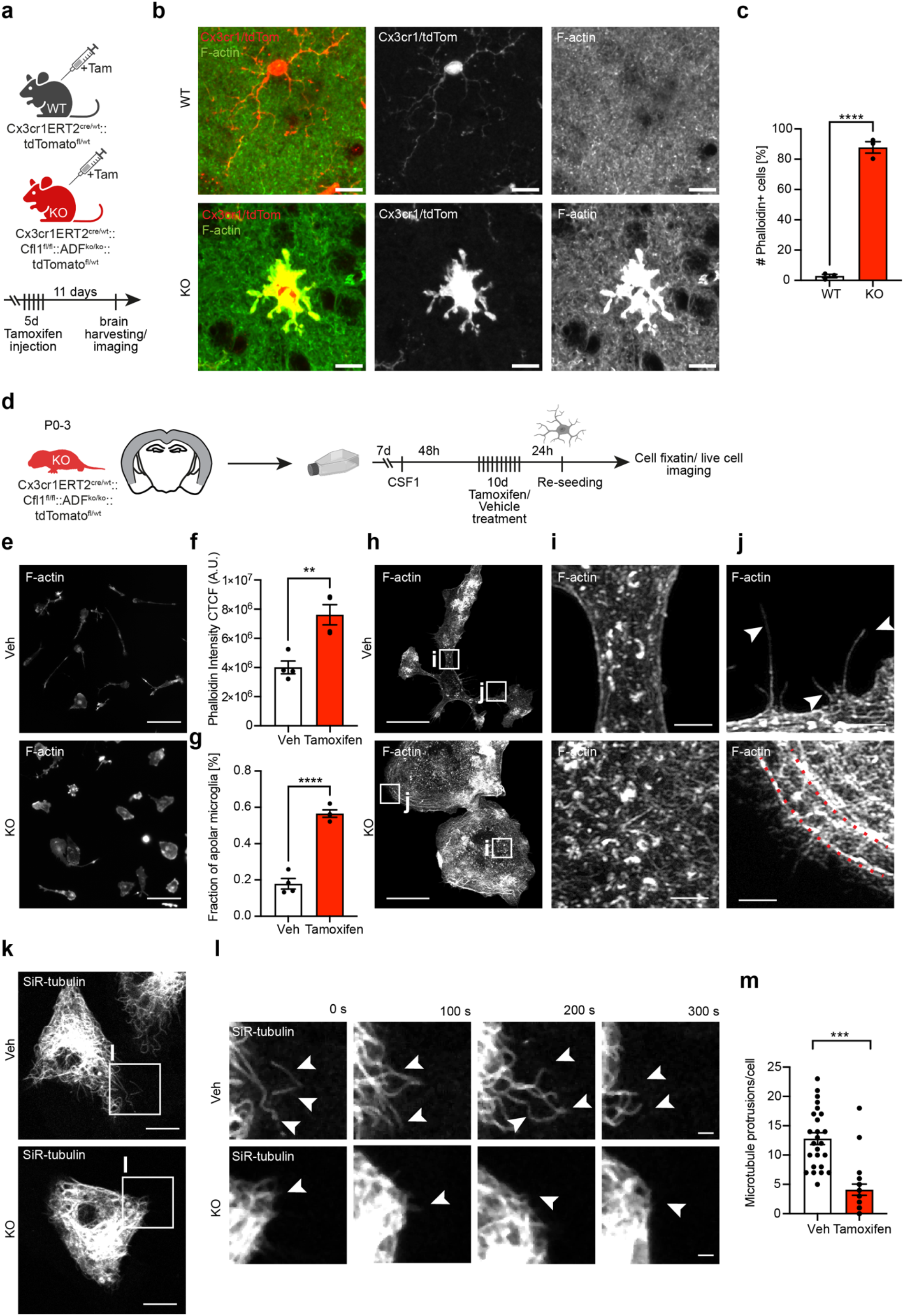
ADF/Cofilin1 deletion profoundly alters the microglial cytoskeleton and cytoskeleton dynamics. **a:** Transgenic mouse lines. WT: Tamoxifen inducible Cx3cr1-dependent tdTomato expression; KO: knock-out of ADF, inducible Cx3cr1-dependent KO of Cfl1 and tdTomato expression. Experimental paradigm with Tamoxifen injection. **b**: Confocal images of microglia with tdTomato expression and Phalloidin staining for F-actin in the somatosensory cortex. Scale bar: 10 µm. **c**: Fraction of microglia with accumulated F-actin in %; two-tailed t-test, t=21.51; df=4, P<0,0001. **d**: Experimental scheme for primary microglia culture preparation from pups. **e**: F-actin staining (Phalloidin) in cultured ADF/Cfl1 microglia treated with vehicle (Veh) control (Methanol) or Tamoxifen. Scale bar: 20 µm. **f**: Phalloidin intensity as corrected total cell fluorescence, CTCF. Unpaired two-tailed t-test: t=4.412, df=6, p=0.0045. **g**: fraction of apolar microglia. Unpaired two-tailed t-test t=10.93, df=6, p=0.000035. **h**: Airyscan super resolution images of phalloidin signal (F-actin) in vehicle and tamoxifen treated primary microglia culture. Scale bar: 20 µm. Intensities have been adjusted to better visualize the F-actin structures. **i, j**: Close ups from **h** for visualization of F-actin puncta (i), F-actin protrusions and accumulation (j). Scale bar: 2 µm. Intensities have been adjusted to better visualize the F-actin structures. **k**: Visualization of tubulin in primary microglia culture by SiR-tubulin labelling. Scale bar: 10 µm. **l**: Close-up images of SiR-tubulin live cell imaging over 5 min at intervals of 100 s. **m**: Number of microtubule protrusions/cell during 5 min. Two-tailed unpaired t-test: t=6.182, df=44, ^***^p<0.0001.

### Absence of ADF/Cfl1 in microglia affects learning and memory

Previous findings from our group and others point to a relationship of microglial fine process motility and neuronal activity^19,26-28^. We hypothesized that our strongly affected microglia in ADF/Cfl1-KO mice might influence neuronal activity and impact cognitive functions like learning and memory. Hence, we carried out behavior experiments to test our hypothesis. First, we investigated explorative behavior in an open field arena. We did not detect any significant changes in the traveled distance between ADF/Cfl1-KO and WT animals **(Fig. 5a, b)**. Next, we performed contextual fear conditioning (cFC) during which mice are encoding a specific context that is subsequently paired with the delivery of received electric foot-shocks. After 48 h mice were placed back into the same context, but without application of foot-shocks to measure freezing levels that are used as a readout for memory retrieval (RE) **(Fig. 5c, d)**.

**Figure 5:**
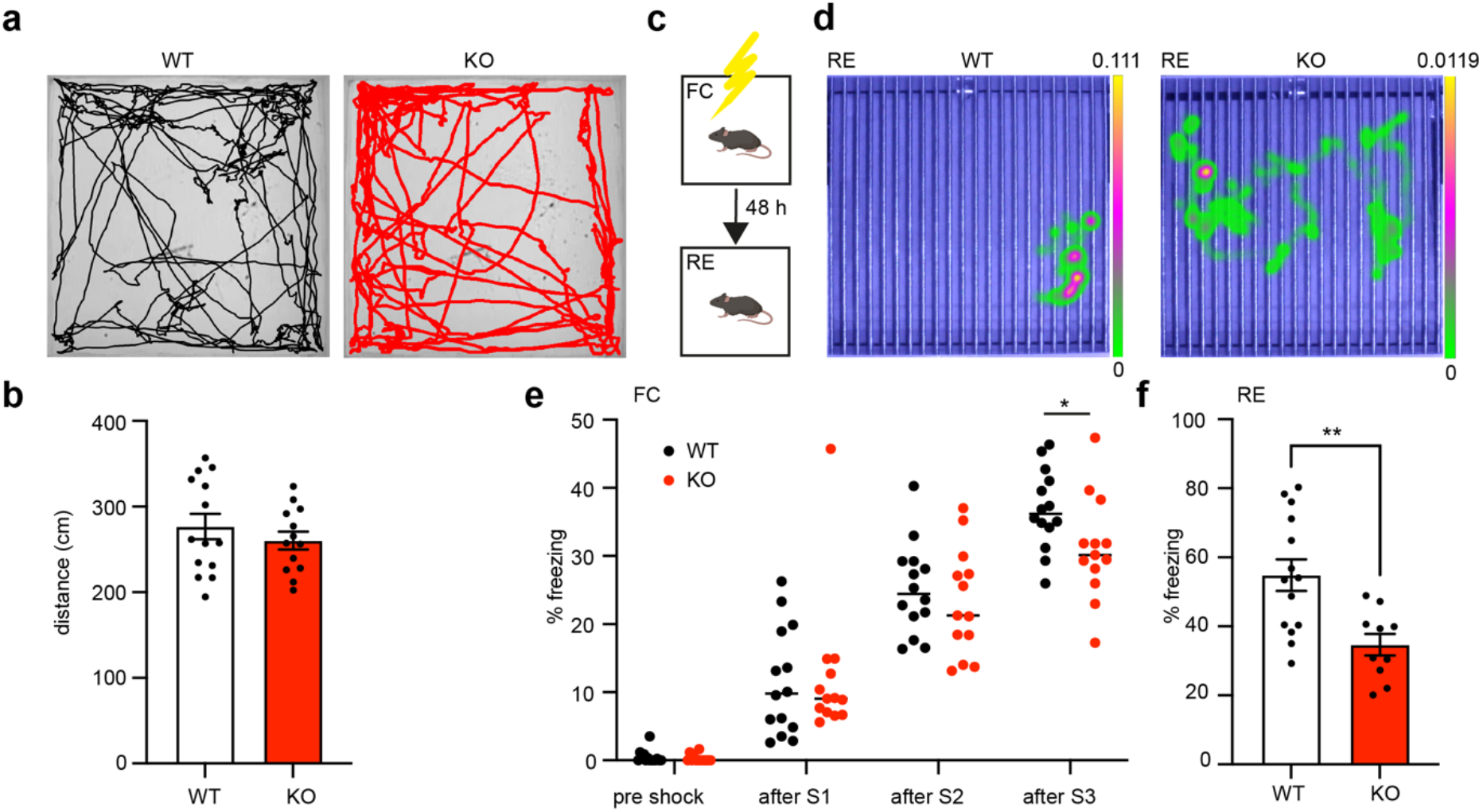
Impaired learning and memory in ADF/Cfl1-KO mice. **a**: Exemplary tracks of open field experiment of WT (black) and KO (red). **b**: Distance travelled of individual mice. WT n=14, KO n=13. Two-tailed t-test, t=0.8826, df=25, p=0.3859. **c**: Experimental design of contextual fear conditioning with fear conditioning (FC) with three foot-shocks and retrieval (RE) after 48 h. **d**: Exemplary heat maps of movement at RE. **e**: Freezing behavior during fear conditioning. Multiple t-tests. Pre shock: t= 0,7088, p= 0,485003, after S1: t= 0,2163, p= 0,830509, after S2: t=0,6764, p= 0,505026, after S3: t= 2,208, p= 0,036619. df=25. WT n=14, KO n=13 mice. **f**: Freezing behavior during RE. Two-tailed t-test t=3.525, df=22, p=0.0031. WT n=14, KO n=10 mice.

During memory encoding, we observed that ADF/Cfl1-KO mice showed significantly less freezing after the third shock in comparison to WT mice **(Fig. 5e)**. In addition, during memory retrieval, ADF/Cfl1-KO mice froze less than WT mice **(Fig. 5f)**. These results point towards impaired memory formation and recall in ADF/Cfl1-KO mice suggesting that an intact actin cytoskeleton might be important for microglia to properly execute their role in cognitive processes like learning and memory.

## Discussion

Here, we show that ADF/Cfl1-deficient microglia are crucially affected in their integrity, morphology and function. The strongly altered morphology prevents microglia to fulfill their function to constantly screen the brain parenchyma. In addition, they lose their capacity to adequately migrate towards a brain injury and initiate shielding of the lesion. These cellular changes are accompanied by increased accumulation of stabilized filamentous actin within microglia due to lack of actin depolymerizing factors ADF/Cfl1. Ultimately, these microglial impairments lead to cognitive deficits.

We generated inducible ADF/Cfl1-KO mice to study the role of these actin depolymerizing proteins in microglia physiology. Manipulations of other individual proteins involved in regulating the actin cytoskeleton in microglia have been shown to affect microglial morphology^20,21,29-31^. In comparison, our double knockout of *Adf* and *Cfl1* has the most drastic effect on morphology, fine process motility and migration. Thus, our findings underscore the crucial position of ADF/Cfl1 for actin filament dynamics. Switching off actin filament depolymerization dramatically changes the morphology of microglia.

Constant surveillance of the brain parenchyma is an elementary physiological feature of microglia^1^. In the absence of ADF/Cfl1, we observed a marked decrease in scanning behavior of microglia in the intact brain. Hence, we show for the first time that ADF/Cfl1 are crucial for cytoskeleton dynamics, thereby revealing a key protein necessary for microglia motility in the healthy brain. Additionally, since ADF/Cfl1 are actin binding proteins, the fundamental implication of actin in microglia motility can be postulated at this point. In other studies, a potential regulatory role of Rac1 in microglial protrusion dynamics and cell motility has been described^20^. Downstream of Rac1, actin modulatory proteins such as Arp2/3 and cyfip1 are involved in microglia surveillance of the brain^30^. This surveillance function of microglia is regulated by cAMP in filopodia via P2Y12 and ATP signaling^3^. It remains to be determined, whether ADF/Cfl1-deficient microglia still sense signaling molecules to a physiological extend but cannot react accordingly due to their restricted motility. In addition, it remains an open question how ADF/Cfl1 is regulated during process motility. An exclusive regulation by p27 and RhoA is unlikely, because p27-KO microglia did not show changes in surveillance behavior ^29^. In line with other studies in neurons, RhoA regulates the actin motor myosin II, but not cofilin during development and in adulthood^32,33^. In addition, ADF/Cfl1 activity is a key factor for enhanced axon regeneration by promoting actin turnover^13^. Together, this further supports our findings that ADF/Cfl1 is crucial for actin filament depolymerization enabling dynamic cytoskeleton rearrangements required for microglial fine process motility.

By performing a laser lesion and monitoring microglia migration in the somatosensory cortex, we found that ADF/Cfl1 is necessary for microglia protrusion extension and migration towards the injury. ATP-release at the lesion is a strong signal to attract and recruit microglia ^2^. Within minutes, microglia extend their protrusions towards the injury site, but only after several hours directed movement and proliferation is initiated^22^. Similar results have been obtained in glioblastoma research, where microglia acquire an ameboid shape and migrate towards injected tumor cells^34,35^. Interestingly, different microglial subsets interact with invading tumor cells in the corpus callosum^36^. Microglia migration has been shown to depend on sex and interferon gamma signaling^23^. We have recently shown that microglia migration is modulated via CX3CR1, potentially via a link to the actin cytoskeleton^24^. Thus, several factors can induce and influence the migration of microglia. Our data complement these findings, demonstrating that ADF/Cfl1 are necessary for microglial migration.

We aimed to further understand the role of ADF/Cfl1 regarding cytoskeletal components in morphology and cellular dynamics. In the absence of ADF/Cfl1, we found accumulated F-actin in microglia in brain sections, as well as in primary microglia, suggesting that ADF/Cfl1 are necessary for actin depolymerization in microglia. Moreover, aberrant actin distribution and decreased cellular protrusions were observed by super resolution microscopy confirming aberrant actin organization in ADF/Cfl1-deficient microglia. In neurons, so called actin arcs, which are located at the growth cone, hinder microtubule protrusion into the leading edge^32,33,37^. We observed similar filamentous actin structures along the border of ADF/Cfl1-deficient microglia, and therefore also investigated the dynamics of tubulin. Live cell imaging of tubulin revealed undirected movement and decreased number of microtubule-directed protrusions indicating a global effect on other components of the cytoskeleton. Interestingly, microtubule dynamics itself have been shown to be essential for microglial function and states^38^. Filamentous actin along the cellular borders/membrane ruffles has been found in activated microglia^39^, suggesting that ADF/Cfl1 regulation might be involved in inducing microglia activation. Hence, targeting ADF/Cfl1 might represent a therapeutic approach for interfering with microglia activation in disease states. Additionally, ADF/Cfl1-deficient primary microglia barely moved within a time course of 3 hours in comparison to vehicle-treated microglia, suggesting impaired capability of spontaneous motion. These results further support our *in vivo* findings that ADF/Cfl1-deficient microglia are less ramified and do not show directed migration. Since we treated ADF^ko/ko^::Cfl1^fl/fl^::Cx3cr1CreERT2^cre/wt^ primary microglia with either vehicle (MeOH) or tamoxifen, we can conclude that ADF-deficiency alone is not sufficient to induce a morphological or motility phenotype. Only the double knockout of *Adf* and *Cfl1* resulted in the prominent changes of microglia morphology, accumulation of stabilized F-actin and decreased tubulin dynamics. Hence, the knockout of both, *Adf* and *Cfl1* is necessary to deplete actin depolymerizing function. In resting microglia, F-actin is mainly found in their cellular processes facilitating efficient surveillance and response to the brain’s environment^3^. Disruption of actin cytoskeleton dynamics by cytoskeletal drugs cytochalasin D and jasplakinolide in primary microglia cultures resulted in reduced microglia migration and proliferation, underscoring the importance of the actin cytoskeleton in those functions^40^. The regulation of actin dynamics by molecules like Rho GTPases, such as RhoA, Rac1, and Cdc42, further underline the complexity of actin-mediated microglial functions. It was shown in RhoA-KO microglia that inactive phosphorylated cofilin was increased and that actin polymerization by staining of F-actin with phalloidin was decreased^31^, which is contradictory to our observation where we can directly link the absence of ADF/Cfl1 to increased F-actin. We can conclude that the aberrant actin filament increase abolishing F-actin polymerization leads to impaired cytoskeletal dynamics, including changes in microtubule dynamics.

We found that ADF/Cfl1-KO mice displayed impaired memory formation and recall. One potential way how microglia might influence cognitive functions, like learning and memory, would be via their interaction with neurons and synapses. Previous studies described a close relationship of microglial fine processes motility and neuronal activity, as well as structural plasticity of synapses. Microglia interaction with synapses in the visual cortex correlated with sensory experience, suggesting that microglia might be involved in experience-dependent modification of synapses in the visual cortex^27^. In addition, noradrenalin which is sensed by microglia in the cortex, changed their screening activity^16,41^. We discovered that microglial screening activity in the hippocampus correlated with the activity of CA1 pyramidal neurons. Indeed, glutamate release in the hippocampus was required for microglial screening and contact of hippocampal synapses^19^. The discovery of synaptic material within microglia suggested that microglia can phagocytose synapses or at least induce their elimination^42^. In addition to elimination, microglia might also mediate the formation of new synapses via BDNF-signaling, since BDNF-deficiency in microglia decreased spine gain in the motor cortex after a motor learning task^17^. These findings underscore that microglia play a role in synapse remodeling and thereby can change the hard wiring of neurons. In this way, microglia can modulate and influence sensation, motor execution, and cognition. It is tempting to speculate that ADF/Cfl1-deficient microglia cannot fulfill this function anymore, which would explain the observed learning and memory deficits. In support of our hypothesis, interfering with the RhoGTPase Rac1 or RhoA, which are both upstream of Cfl1, decreased the performance of mice in a novel object recognition task^20^. Thus, our results corroborate previous findings that an intact actin cytoskeleton in microglia is a prerequisite to fulfill their synapse remodeling function, thereby supporting higher cognitive functions like learning and memory.

## RESOURCE AVAILABILILTY

### Lead contact

Further information and requests for resources and reagents should be directed to and will be fulfilled by the lead contact, Martin Fuhrmann.

### Materials availability

All unique reagents generated in this study are available from the lead contact with a completed material transfer agreement.

### Data and code availability

The authors declare that the data supporting the findings of this study are available within the paper, the methods section, and Supplementary data files. Data are available open access from DRYAD.

## ACKNOWLEDGEMENTS

This work was supported by the DZNE, and grants to MF by the European Union ERC-CoG (MicroSynCom 865618). MF, SC received funding from the German research foundation DFG (SFB1089 C01, B06; SPP2395) and MF is a member of the DFG excellence cluster ImmunoSensation2. This work was also supported by the iBehave and CANTAR network to MF, FN, SP (funded by the Ministry of Culture and Science of the State of North Rhine-Westphalia; the funders had no role in study design, data collection, and interpretation, or the decision to submit the work for publication). FN received funding from the Mildred-Scheel School of Oncology Cologne-Bonn. We would like to thank the Core Research Facilities, especially the Light microscopy facility (LMF), Services at DZNE, and the DZNE Animal Research Facilities for continuous support through the project. Figures were prepared with Illustrator CS5 Version 15.0.1 (Adobe) and created with with BioRender.com (Details see below)

## AUTHOR CONTRIBUTIONS

SC, MDR, SP, FCN and MF conceptualized the experiments, collected data, analyzed data, designed the figures and wrote the manuscript. JS carried out the experiments on primary microglia, analyzed data and prepared figures. MM, FM carried out experiments and analysed data. SJ, KMW, AB helped with histology, genotyping and breeding. CG generated ADF/Cfl1-KO mice. WW provided resources and consulting on conceptualization. FB provided resources, supervised the cell culture experiments and provided conceptualization input. MF together with SC supervised the project. MF provided resources, provided technical expertise, analyzed data, conceptualized experiments, wrote the manuscript.

## Methods

### Mice

Mice were group-housed with a maximum of five animals per cage and separated by gender. The animals were subjected to a day/night cycle of 12 h and were able to access water and food supply *ad libitum*. All experiments were performed according to animal guidelines and were approved by the “Landesamt für Natur, Umwelt und Verbraucherschutz (LANUV)” of North-Rhine Westphalia (Office for Nature, Environment and Consumer Protection of North Rhine-Westphalia) in Recklinghausen, Germany, under the licenses Az84-02.04.2017.A098, Az81-02.04.2023.A102, Az2024-144. We generated a quadruple transgenic mouse line ADF ^KO/KO^::Cfl1^fl/fl^::Cx3Cr1CreERT2^cre/wt^::Gt(Rosa26)tdTomato^fl/fl^, based on Cx3Cr1-CreER: B6J.B6N(Cg)-Cx3Cr1tm2.1(cre/ERT2)Jung/J (Jackson Laboratory), and Gt(Rosa)26-tdTomatofl: B6.Cg-Gt(ROSA)26SORtm14(CAG-tdTomato)HZE/J (Jackson Laboratory. The ADF/Cfl1 mouse line was kindly provided by Walter Witke (University of Bonn, Germany)^8^. Our transgenic mouse line carries an *Adf* knockout together with an inducible conditional knockout of *Cfl1* in *Cx3cr1*-expressing cells. Tamoxifen application leads to Cre-recombinase expression in CX3CR1^+^ microglia, which recombines the loxP-flanked alleles *Cfl1* and the fluorescence reporter *tdTomato*. The mouse line Cx3Cr1CreERT2^cre/wt^::Gt(ROSA)tdTomato^fl/fl^ served as a control. Genotyping PCRs were performed using mutant-specific, custom-made primers (Supplementary information).

### Tamoxifen injections in vivo

To induce Cre-recombinase expression under *Cx3cr1*-promoter, adult mice were injected intraperitoneally (i.p.) with 0.1 mg/g body weight (BW) Tamoxifen (Sigma-Aldrich) for five consecutive days. Tamoxifen was dissolved in Miglyol (Miglyol 812 Hüls Neutralöl; Caesar & Loretz) using sonication and was delivered in a volume of 5 µl/g body weight.

### Primary microglia culture

Mixed glial cultures were prepared from 0 – 3 day old mice according to the dissection procedures previously described^43^. Cerebral cortices from P0–3 mice were isolated, meninges were carefully removed, the cortices were subsequently mechanically dissociated in Hanks’ Balanced Salt Solution (HBSS; Life Technologies) and trypsinized in 0.05% trypsin (Life Technologies) for 15 min at 37 °C. Afterwards, trypsin was inhibited by triturating cells in complete Dulbecco’s modified Eagle’s medium (Life Technologies) supplemented with 1% Penicillin/Streptomycin (Gibco) and 10% heat-inactivated FBS (PAN-Biotech). Cell suspensions were subsequently plated into T75 flasks, coated with collagen-I (0.025%, Sigma) and 100 µg/ml poly-ornithine (Sigma). Cells were grown at 37°C and 5% CO2 in a humidified atmosphere until 90% confluency (usually within 7 days). Subsequently, microglia proliferation was increased by treatment with 100 ng/ml murine Colony-stimulating factor-1 (M-CSF1) (Peprotech) for 48 hours.

### Hydroxytamoxifen treatment in vitro

Induction of Cre-recombinase *in vitro* was done 48 hours after the first M-CSF-1 stimulation. Mixed glial cultures were treated either with 1 µM (Z)-4-Hydroxytamoxifen (Sigma) dissolved in Methanol (MeOH) or vehicle control (corresponding amounts of MeOH) for 10 days. For the first 5 days, the medium was replaced daily by fresh medium, containing 1 µM hydroxytamoxifen or MeOH. On the 5^th^ day of treatment, new CSF-1 (100 ng/ml) was added to the hydroxytamoxifen- or MeOH-containing medium and cells were left to grow for additional 5 days. Then, microglia were harvested by rigorously tapping the flask. The supernatant containing the floating microglia was centrifuged for 5 min at 150 xg. The cell pellet was resuspended in culture medium (DMEM, 10% FCS, 1% P/S) and plated on collagen-I (0.025%, Sigma) and 100 µg/ml poly-ornithine (Sigma)-coated plastic dishes for RNA and protein analysis (Thermo Fisher), or collagen-I (0.025% Sigma) and 100 µg/ml poly-ornithine-coated (Sigma) glass coverslips for immunocytochemistry or live cell imaging dishes (4 well µ-slides dishes, IBIDI) for SiR-actin and SiR-tubulin imaging.

### Cortical cranial window preparation

Cortical cranial window surgery was performed in adult mice and four weeks prior to *in vivo* imaging. Mice received buprenorphine (i.p., 0.05 mg/kg BW) 20 min before surgery and were subsequently anaesthetized by administration of Ketamin/Xylazin (i.p., 0.13/ 0.01 mg/g BW). After reaching surgical tolerance, the mouse was placed in a stereotactic frame, while body temperature was maintained at 37°C by a heating pad. Subcutaneous application of Dexamethasone (s.c., 0.2 mg/kg BW) and Carprofen (5 mg/kg BW) was applied shortly prior to surgery to prevent swelling of the brain. Under surgical conditions, the skin on top of the skull was removed with surgical scissors, and a circular piece (Ø 4 mm) of the skull above the somatosensory cortex was removed using a dental drill. The dura was carefully segregated, and the brain surface was rinsed with sterile saline subsequently. For the window, a 4 mm diameter glass coverslip was inserted into the craniotomy and glued to the skull-bone with dental cement. A headpost (Luigs & Neumann) for head-fixation during *in vivo* imaging was mounted alongside the imaging window. For post-surgery analgesia, buprenorphine (s.c., 0.05 mg/kg BW) was applied every 8-12 h for 3 days.

### In vivo two-photon imaging

Mice were anaesthetized with Isoflurane (0.9 % v/v in oxygen) and head-fixed under the microscope, while body temperature was maintained at 37°C by a heating pad.

Microglial fine processes motility recording: Imaging of microglial process motility was performed 11 days after last tamoxifen injection. Z-stacks were recorded every 5 min with a Zeiss LSM 7MP microscope (ZEISS) equipped with a Ti:Sapphire laser (Chameleon Ultra II, Coherent). Excitation wavelength was tuned to λex=980 nm and a ZEISS 20x water immersion objective (W Plan-Apochromat 20x/ NA1.0 water immersion). Red tdTomato fluorescence was bandpass filtered (BP590/40) and collected on a GaSP-detector. Image volumes were acquired at a depth of 150 µm with 106 µm x 106 µm size in x, y-direction and 70 µm in axial z-direction. A z-spacing of 1 µm was used and a pixel size of 0.104 µm/pixel. Six volumes were recorded with 5 min time-intervals (start to start). For density quantification of microglia, we recorded a volume spanning 300 µm from the brain surface. The x,y-dimensions were 607 µm x 607 µm and the z-step lengths was 3 µm. The pixel size was 0.4 µm/pixel.

To record microglia migration towards a laser-induced lesion, the tissue injury was induced at a depth of 150 µm by setting the area to 10 x 10 µm^2^ in x, y-direction, with 0.04 µm/pixel, λex=800 nm, 100 % laser power and 6.3 µs/pixel dwell time for 30 s. 3D-time-lapse acquisition started 5 min before the lesion induction (0h) and continued 8 hours later with 4-hour time-intervals up to 20 hours post lesion. At every time point, volumes with 607 x 607 x 300 µm^3^ in x, y, z-dimension starting from brain surface, were acquired with the Zeiss LSM 7MP microscope. The z-step distance was 3 µm.

### Life cell imaging

SiR-Tubulin and the efflux pump inhibitor verapamil were purchased as a kit (Spirochrome). Labeling of microglia was performed modifying the manufacturer’s recommendations: For labeling with SiR-Actin, cells were treated with 10 µM verapamil and 1 µM SiR-Actin by adding both components directly to the medium and incubating at 37°C and 5% CO2 for 3 h. For SiR-Tubulin labeling, cells were treated with 1 µM SiR-Tubulin and 10 µM verapamil for 1 h at 37°C and 5% CO2 prior to imaging. The medium was kept on the cells during the imaging. Imaging was performed with the Zeiss LSM980 and a 40x oil objective at 37°C, 5% CO2 and a humidified atmosphere.

### Histology

*Ex vivo* tissue: Mice were deeply anesthetized by an i.p. injection of overdosed Ketamin/Xylazin (Medistar/Bayer) and transcardially perfused with phosphate-buffered saline (PBS). The brains were removed and fixed in 4% (w/v) Paraformaldehyde (Carl Roth) overnight. 100 µm thick brain slices were cut on a vibratome (Leica VT 1200S). For antibody staining, slices were incubated overnight at 4°C free-floating in blocking buffer (0.4% Triton X-100 (Sigma), which contained 4% normal goat serum (Invitrogen), 4% bovine serum albumin (BSA; Sigma) in PBS) and primary antibody rabbit anti-Iba1 (1:1000; FUJIFILM Wako Pure Chemical Corporation, 019-19741). After three washing steps with PBS, secondary antibody goat anti-rabbit Alexa Fluor® 488 (1:400; Invitrogen, A11008) was applied for 2 h at room temperature. Afterwards, slices were washed three times in PBS, mounted onto glass microscopy slides using fluorescent mounting medium (Agilent), and covered with a glass coverslip.

For F-actin staining, brain slices were incubated in blocking buffer, containing Alexa Fluor® 488 Phalloidin (1:400; Invitrogen, A12379) for 2 h at room temperature. Afterwards, three washing steps with PBS were performed, slices were mounted onto glass microscopy slides using fluorescent mounting medium (Agilent) and covered with a glass coverslip.

#### In vitro primary cell culture

Primary microglia were fixed for 20 min at room temperature in PFA-PHEM-sucrose (4% Paraformaldehyde (Carl Roth) in PHEM buffer ((25mM Hepes, 60mM Pipes, 10mM EGTA, 2mM MgCl2), supplemented with 4% sucrose (Carl Roth); the pH was adjusted to 7.4) to optimally preserve the cytoskeleton. Subsequently, cells were washed three times in PHEM buffer and permeabilized for 3 minutes in 0.02% Triton-X 100 (Sigma). Cells were then blocked in 5% BSA (Sigma) in PHEM buffer for 1 h at room temperature. For labelling of F-actin, Alexa Fluor® 488 Phalloidin (1:1000; Invitrogen, A12379) was added to PHEM buffer and incubated for one hour at room temperature protected from light. Subsequently, cells were washed three times with PHEM buffer and mounted on objective slides in aqueous mounting medium (Fluoromount, Sigma).

### Confocal imaging

For confocal imaging of brain slices to measure microglia morphology and phalloidin signal, a LSM Zeiss 700 upright was used with an 20x objective (Plan-APOCHROMAT 20x / 0.8 NA). Phalloidin and tdTomato fluorescence were excited at 488/555 nm with an argon respectively HeNe-laser. Emitted fluorescence of Phalloidin and tdTomato was bandpass filtered (BP500-550; BP580/30), the pinhole was set to an Airy unit of 1, the pixel size was 0.31µm/pixel, the image size was 640 x 640 µm^2^, and the pixel dwell time 0.64µs/pixel. For imaging of primary microglia, a Zeiss confocal microscope (LSM980), optionally with the Airyscan super resolution, or a Zeiss epifluorescence microscope (EpiScope2), was used.

### Open field

To investigate the explorative behavior of ADF/Cfl1-KO in comparison to control mice, an open field behavior experiment was performed. For this, mice were placed in a 50 x 50 cm box for 5 min, individually. Movement of mice during exploration behavior was recorded using a video camera and analyzed using EthoVision XT (Noldus).

### Contextual fear conditioning

Fear conditioning (FC) was performed in a custom-made cage with a steel-grid floor. Mice were allowed to explore the cage for 120 s. Three electric foot shocks were applied (0.75 mA for 2 s) with an inter-shock interval of 60 s. Retrieval (RE) was performed 2 days after fear conditioning. The animal’s movement was recorded using a video camera and analyzed semi-automatically with EthoVision XT (Noldus).

### Image data analysis

#### Analysis of cell morphology ex vivo

To determine the soma diameter, maximum intensity projections of z-stack images were generated in Fiji, followed by manual diameter measurement. In each field of view (FOV), 9-10 cells were analyzed, with three FOVs per animal. The mean of each FOV was determined, averaged over animals and plotted using GraphPad Prism. Microglia cells (Iba1) were 3D reconstructed in IMARIS (Oxford Instruments) using the “filament tracing” tool. 3D Sholl analysis was carried out in IMARIS.

#### Microglial motility analysis

In general, microglial motility was analyzed as described before^19,44^. Briefly, z-stacks were initially rigidly registered using subpixel image registration by cross-correlation ^45^ provided by the open-source Python image processing library Scikit-image^46^. Individual stacks were then median filtered and individual microglia were identified by scrolling through the stacks at each time point and cropped using ImageJ. Z-stacks spanning 35 μm in depth were isolated from the original z-stack. Stacks were registered by applying the ImageJ ‘StackReg’ plugin^47^. For all time points, average intensity projections were created resulting in a 2D visualization of microglia and their fine processes. Individual 2D images were merged into a time-lapse. Time points were pseudo colored in magenta, cyan and grey. Magenta areas account for lost, cyan for gained and grey for stable microglial fine processes.

The percent gained, lost and stable areas were calculated by dividing the number of gained, lost and stable pixels by the number of all pixels (sum of gain, loss and stable). The turnover rate (TOR) of individual microglia processes was calculated as the number (absolute pixel value) of lost, Nlost (red), and newly gained, Ngained (green) pixels divided by the sum of all pixels within a determined region of interest (ROI).

We used a custom analysis pipeline for automated quantification ot the turnover rate of microglial protrusions, written in Python as described before ^48^. We calculated the temporal variation *ΔB(ti)* of binarized images by subtracting the binarized image *B(ti+1)* at time point *ti+1* from the binarized image *B(ti)* at time point *ti*: *ΔB(ti) = 2×B(ti+1)-B(ti)* for *i=0, 1, 2, …, N-1*, where *N* is the total number of all time lapse time points. Pixels in *ΔB(ti)*, that have the value 1, were categorized as stable pixels, whereas pixels with the value -1 were categorized as gained pixels, and pixels with the value 2 as lost pixels. The microglial fine process motility was then assessed by calculating the turnover rate (TOR) as the ratio of the number of all gained pixels *Ng(ti)* and all lost pixels *Nl(ti)* divided by the sum of all pixels: *TOR(ti) = (Ng(ti)+ Nl(ti)) / (Ns(ti) + Ng(ti)+ Nl(ti))*, where *Ns(ti)* is the number of all stable pixels. The average turnover rate 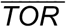 was calculated by averaging *TOR(ti)* over all *N-1* time lapse time

#### Microglia migration

Microglia migration was analyzed with FIJI “manual tracking” plugin. After acquisition the images were aligned in z,t-dimension by manually aligning the top and bottom image for each time-point. 3D-registration was carried out using the FIJI “TurboReg” plugin. Subsequently, image stacks were median filtered and twenty slices, spanning 60 µm and containing the lesion, were maximum intensity projected. These images were exported to Fiji and aligned in x,y-dimension over the whole imaging time-period using the “StackReg” plugin. Individual microglia were tracked over the imaging time-period using the FIJI “manual tracking” plugin. Two circles were drawn around the lesion center with a radius of 140 and 250 µm. The number of microglia per each circle segment (0-140 µm, 140-250 µm and >250 µm) was measured for each time-point and mouse. Data were normalized to the time-point 0h. For all microglia that translocated more than 10 µm over the 20h recording period the directionality index DI, the Euclidian traveled distance T and the perfect distance traveled (L_1_) were calculated. DI was calculated by dividing the difference in Euclidian distance to the lesion (L1) by the difference in Euclidian distance of the traveled distance (T) **(Fig. 3i)**. Therefore, we measured the position (x,y-coordinates) of each microglia at each time-point (t0h – t20h) and the position of the lesion center. The total Euclidian distances (L) and (L2) between the positions at t0h and t20h in relation to the lesion center were calculated for each cell. In addition, the Euclidian distance (T) of the traveled distance was calculated between position t0h and t20h for each cell. The perfect distance traveled in direction to the lesion (L1) was calculated by subtracting L2 from L.

#### Cell density in vivo

Microglia density was measured in a volume of interest with 400 x 400 x 63 µm^3^ dimension, 100 µm beneath the brain surface, using the spot detection by IMARIS (Oxford Instruments).

#### F-actin analysis ex vivo and in vitro

Microglia with accumulated F-actin in cortical brain slices were counted with the “CellCounter” plugin in FIJI. The percentage of double positive (Phalloidin & tdTomato) microglia in relation to all Cx3cr1-tdTomato positive cells was determined by manually counting in 300 x 300 µm^2^ large field of views (FOV). At least three FOVs were analyzed per mouse and average values were calculated for each mouse. The data were plotted and analyzed with GraphPad Prism. To quantify F-actin content in primary microglia cells *in vitro*, the intensity of the phalloidin labeling was compared between images. All images were acquired with the same settings and within one imaging session using an epifluorescence microscope (Zeiss, Episcope2). Fluorescence intensity was analyzed by outlining the cells with the freehand selection tool. Using the ‘Measure’ function in Fiji, the cell area, Integrated density and Mean Gray area were measured. One background intensity measurement was performed for each image using the rectangle tool. Subsequently, the CTCF ((CTCF) = Integrated Density – (Area of Selected Cell x Mean Fluorescence of Background values) was calculated for each cell. The experiment was performed with n=4 mice, analyzing a minimum of 28 cells for each. Results were averaged for each biological replicate, and data were plotted using GraphPad Prism. A two-tailed unpaired t-test was performed for statistical analysis.

#### Cell morphology in vitro

To assess whether cells displayed a polar or apolar morphology, a trained scorer manually classified cells as polar or apolar. Criteria for polarity included detectable branches or a clear orientation/elongation of the cell. Cells were counted using the “CellCounter” Plugin in Fiji. The percentage of apolar cells was calculated by dividing the number of apolar cells through the number of total cells. Data were plotted in GraphPad Prism, and a two-tailed unpaired t-test was performed for statistical analysis.

#### Microtubule dynamics in vitro

The number of microtubule protrusions per cell were manually counted in Fiji. For each video, the first frame was analyzed. Microtubule protrusions were counted for each cell manually. A protrusion was classified as a microtubule protruding into the periphery of the cell and out of the dense network of microtubules present in the cell body. A minimum of 21 cells were analyzed per experimental condition. Results were plotted in GraphPad Prism and a two-tailed unpaired t-test was performed for statistical analysis.

## Statistics

Quantifications, statistical analysis, and graph preparation were carried out using GraphPad Prism 9 (GraphPad Software Inc, La Jolla, CA, USA). To test for normal distribution of data, D’Agostino and Pearson omnibus normality test was used for sample sizes of n>6 and the Shapiro-Wilk normality test for n<6. To test statistical significance for groups of two normally distributed data sets paired or unpaired two-tailed Student’s t-tests were applied. If data was not normally distributed, Mann-Whitney test was used to compare two groups. One-way ANOVA with Tukey’s or Bonferroni’s multiple comparison test were performed on data sets larger than two, if normally distributed. For comparison of more than two not normally distributed data sets the Kruskal-Wallis test was performed with Dunn’s correction for multiple comparisons. If not indicated differently, data are represented as mean ± SEM.

## BioRender Licenses

Figure1: https://biorender.com/d98s625

Figure2: https://biorender.com/x61v825

Figure3: https://biorender.com/y27g673

Figure4: https://biorender.com/q45r662

Figure5: https://biorender.com/c66o523

FigureS1: https://biorender.com/l06z149

FigureS2: https://biorender.com/e38o359

