## Supplementary information for "Deficiency of actin depolymerizing factors ADF/Cfl1 in microglia decreases motility and impairs memory"

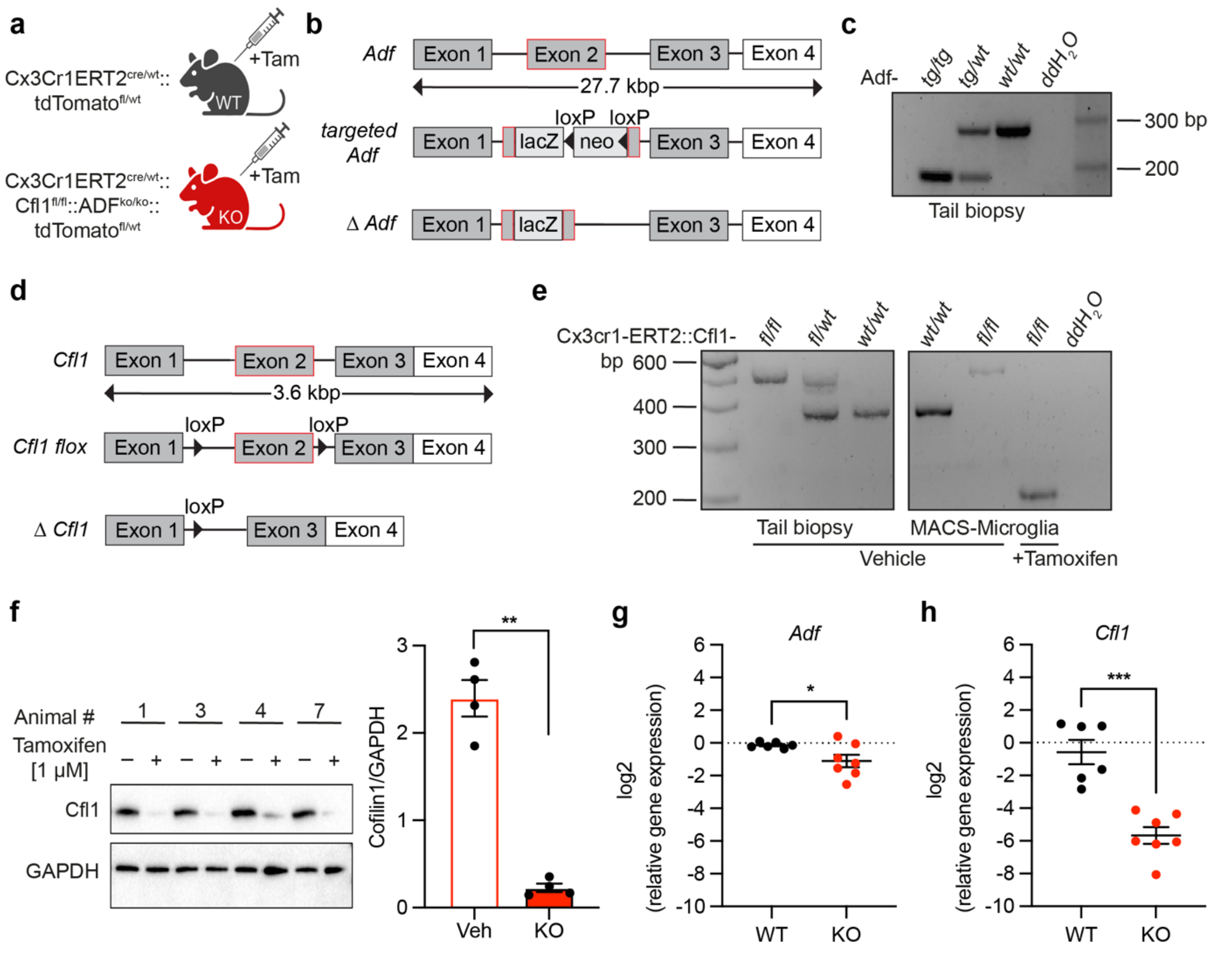


**Supplementary figure 1: Characterization of ADF/Cfl1-KO mice.**

**a**: Schematic overview of mouse lines and respective genotype used as KO and WT, after induction with Tamoxifen, in all experiments.

**b**: Genetic background of ubiquitous ADF knockout, achieved by insertion of a lacZ-cassette into exon 2.

**c**: Verification of ADF knockout on genomic level by PCR. Agarose gel electrophoresis reveals knockout of ADF by a 200 bp PCR product.

**d**: Genetic background for conditional Cfl1 knockout. In the cofilin-1 gene, exon 2 is flanked by two loxP sites that can be cut out by cre recombinase.

**e**: Verification of Cfl1 knockout on genomic level by PCR in tail biopsy and MACS-microglia. Agarose gel electrophoresis of PCR products show a 400bp WT-band and a 500bp Cfl1$^{flox}$-band in tail biopsies respectively microglia from vehicle-treated mice. Microglia from tamoxifen-treated homozygous Cfl1$^{fl/fl}$-mice show an Exon2-deleted 200 bp sized band.

**f**: Verification of Cfl1 knockout on protein level via Western blot in primary ADF-KO/Cfl1fl microglia, treated with Tamoxifen or vehicle. Tamoxifen treated microglia show a statistically decreased Cfl1 protein concentration. Two-tailed t-test (Welch), n= 4, **: p = 0.0013.

**g**: Gene expression level of *Adf* verifies knockout of *Adf* by qPRC on transcriptional level in primary microglia. Two-tailed t-test, n= 6, *: p= 0.0423.

**h**: Gene expression level of *Cfl1* verifies knockout of *Cfl1* in primary microglia after Tamoxifen treatment. Two-tailed t-test, n= 6, ***: p= 0.0001.

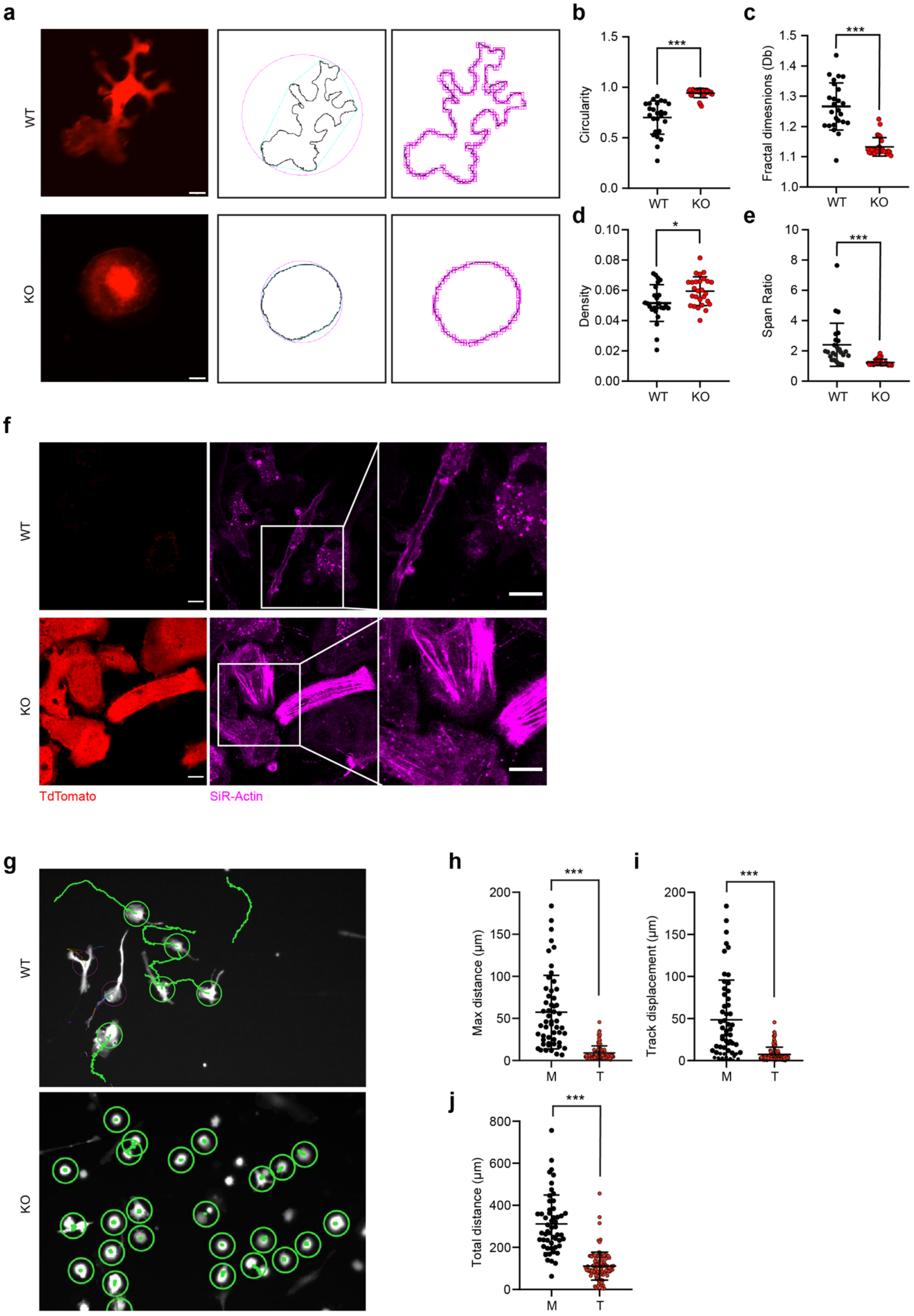
a
WT
KO
b
Circularity
1.5
1.0
0.5
0.0
***
WT
KO
c
Fractal dimesnions (Db)
1.5
1.4
1.3
1.2
1.1
1.0
***
WT
KO
d
Density
0.10
0.08
0.06
0.04
0.02
0.00
*
WT
KO
e
Span Ratio
10
8
6
4
2
0
***
WT
KO
f
WT
KO
TdTomato
SiR-Actin
g
WT
KO
h
Max distance (μm)
200
150
100
50
0
***
M
T
i
Track displacement (μm)
200
150
100
50
0
***
M
T
j
Total distance (μm)
800
600
400
200
0
***
M
T

**Supplemantary Figure 2: Analysis of primary microglia.**
**a**: Microglia expressing tdTomato, treated with vehicle control (Methanol) or Tamoxifen. Scale bar: 10 µm. Example images of convex hull and box counting generated with the FracLac Fiji plugin. **b**: Quantification of circularity (F-test F=13.58, p= 0.0000000045), unpaired two-tailed t-test with Welch's correction (t=7.124629909, df=27.26845182, p= 0.0000001098), **c**: fractal dimensions (Db) (F-test F= 6.602,p= 0.00000931), unpaired two-tailed t-test with Welch's correction (t= t=8.0071842, df=30.647181, p= 0.0000000053).**d**: density (F-test F= 1.639, p= 0.0115), unpaired t-test (t=2.624, df=50, p=0.0115), **e**: span ratio (F-test F= 49.4242285393244, p<0.00000000001), unpaired t-test with Welch's correction (t t=4.05, df=24.89, p= 0.00043).
**f:** Sir-Actin staining of primary microglia comparing WT and KO. Note that WT microglia display actin puncta and few actin filaments, whereas KO microglia contain high levels of filamentous actin. Scale bars: 2 µm.
**g:** Example images of tdTomato expressing microglia treated either with vehicle control (MeOH) or tamoxifen, analyzed with the TrackMate plugin for Fiji. Only green trajectories were included in the analysis.
**h-j:** Average maximum travelled distance, track displacement and total traveled distance for all analyzed primary microglia. All three parameters are significantly reduced comparing WT to KO microglia. (h-j: unpaired t-test with Welch's correction h: 8.011, df=53.87, i: t=6.323, df=53.72, j: t=10.8, df=64.3, *** p<0.0001, WT n= 53, KO N=107 cells)

**Supplementary Methods**

- PCR and primer specifications
- MACS
- SiR Actin
- Trackmate migration (imaging + Analysis)

***Genotyping***

Genotyping of mice was done using the following primers: *Adf* (Sigma) A: 5'-GATTAAGTTGGGTAACGCC-3', B: 5'-GAAGAAGGCAAAGAGATCTT-3', C: 5'-CCCAACATTGCCAATACCATT-3'; *Cfl1* (Sigma) A: 5'-CGCTGGACCAGAGCACGCGGCATC-3', B: 5'-CTGGAAGGGTTGTTACAACCCTGG-3', C: 5'-CATGAAGGTCGCAAGTCCTCAAC-3'; *Cx3cr1* (Eurofins) A: 5'-GTCTTCACGTTCGGTCTGGT-3', B: 5'-CCCAGACAGTCGTTGGTCCTT-3', C: 5' -CTCCCCCTGAACCTGAAAC- 3'; *ERT2* (Sigma) A: 5'-TGAGCCCCCCATACTCTATTCC-3', B: 5'-ATCTCCACCATGCCCTCTACAC-3'; *tdTomato* (Eurofins) A: 5'-AAGGGAGCTGCAGTGGAGTA-3', B: 5'-CCGAAAATCTGTGGGAAGTC-3', C: 5'-GGCATTAAAGCAGCGTATCC-3', D: 5'-CTGTTCCTGTACGGCATGG-3'. *ADF*$^{ko/ko}$ yield a single ~180 bp fragment, *ADF*$^{ko/wt}$ yield a

~180 bp and a ~257 bp fragment, and *ADF*$^{wt/wt}$ a single ~257 bp fragment. *Cfl1*$^{fl/fl}$ yield a single ~ 420 bp prior recombination by cre-recombinase, and a single ~170 bp fragment after deletion of exon 2. *Cfl1*$^{fl/wt}$ yield a ~ 420 bp and a ~ 380 bp fragment, *Cfl1*$^{wt/wt}$ yield a single ~ 380 bp fragment. *Cx3cr1-creERT2*$^{cre/cre}$ yield a single ~ 230 bp fragment, *Cx3cr1-creERT2*$^{cre/wt}$ yield a ~ 230 bp and a ~ 151 bp fragment, and *Cx3cr1-creERT2*$^{wt/wt}$ yield a single ~ 151 bp fragment. *tdTomato*$^{fl/fl}$ yield a single ~ 196 bp fragment, *tdTomato*$^{fl/wt}$ yield a ~ 196 bp and a ~ 297 bp fragment, *tdTomato*$^{wt/wt}$ yield a single ~ 297 bp fragment.

***Western blot***

Cultured microglia were washed once with cold PBS and then lysed in cold radioimmunoprecipitation assay buffer (50 mM TrisHCl, pH 7.4, 150 mM NaCl, 1% Triton 0.5% sodium deoxylcholate, 0.1% SDS, 1 mM EDTA, ddH2O), supplemented with protease inhibitors (Merck, Calbiochem set III) for 20 minutes on ice. Cell lysates were collected in Eppendorf tubes and centrifuged at 20.000 × g. Supernatant was transferred to a tube containing Roti load I SDS sample buffer (Carl Roth). The quantification of protein amount per sample was hindered by strong expression of tdTomato in tamoxifen treated cells, which made measurements obtained from bicinchoninic acid assay-based protein quantification unusable, as this assay is based on photometric evaluation at 562 nm. Instead, the same number of cells was plated in each well and equal volumes of the sample were loaded on 4-12% NuPage Bis-Tris gels (Thermo Fisher) and run at 120V with 1X NuPAGE MOPS running buffer (Thermo Fisher). A Ponceau S (Sigma) stain was performed to verify that equal amounts of proteins were loaded after Western blotting. Western blot analysis was performed by transferring proteins to nitrocellulose or PVDF membranes (activated with 100% Methanol for 1 minute), using a wet blot tank system (Bio-Rad) for 2 h with 1X Transfer buffer (25 mM Tris, 192 mM glycine, 20% methanol). The membranes were then blocked for 1 h at room temperature with 5% skim milk (Applichem) in Tris-buffered saline with 1% Tween20 (TBS-T) before incubating with primary antibodies overnight. Cofilin-1 (Cell signaling, 1:1000) was incubated in 5% Bovine serum albumin (Sigma) in TBS-T and GAPDH (abcam, 1:1000) was incubated in 5% skim milk. Membranes were then washed three times for 10 min each, in TBS-T and incubated with HRP-coupled secondary antibody (Donkey-anti-Rabbit-HRP, Thermo Scientific, 1:3000; Goat-anti mouse-HRP, Merck, 1.3000) for 1 hour, followed by three times of 10 min washes in TBS-T. Blots were incubated with luminol-based HRP substrate developing solution (Thermo Scientific) for 1 minute and membranes were imaged using the a BioRad ChemiDOC imaging system. Quantification of band densities was performed using FIJI. For relative quantifications, measurements were normalized to loading control.

***Gene expression***

The expression of *Adf*, *Cfl1* and was analysed by reverse transcription-quantitative polymerase chain reaction (RT-qPCR), using the PowerUp SYBR green assay (Applied Biosystems). RNA isolation from primary microglia was performed with the RNeasy Mini kit (Qiagen) according to the manufacturer's protocol. RNA yield and integrity was assessed by the TapeStation RNA screentape analysis (Agilent), following the manufacturer's protocol. The concentration of input RNA for reverse transcription was normalized. Afterwards RNA was transcribed into complementary DNA (cDNA) with the iScript cDNA synthesis kit (BioRad) according to the manufacturer.

The RT-qPCR SYBR Green assay was validated in terms of optimal cDNA concentration (dilution series 0.16-20 ng/µl), optimal primer concentration (concentration matrix 0.5-8 µM), optimal primer combination (lowest $C_q$, absence of primer dimers by gel electrophoresis), primer specificity (melt curve), and primer efficiency (standard curve). Suitable and stable reference genes were determined by testing primer candidates with WT, and ADF/Cfl1-KO samples. These reference gene candidates were verified by the web-based tool RefFinder (http://blooge.cn/RefFinder/), which combines the major computational tools for determining the most stable reference gene: geNorm, Normfinder, BestKeeper, and the comparative Δ-Ct method. Reference gene candidates tested were: *Cx3cr1*, *Csf1r*, *Hexb*, *Hprt*. Most stable reference genes for *in vitro* primary microglia were determined to be *Csf1r* and *Cx3cr1*. RT-qPCR was performed in technical duplicates using the StepOnePlus instrument (Applied Biosystems), performing 40 cycles and a melting curve. From resulting Ct values, dCt and relative quantity (RQ) were calculated for reference genes and target genes. Afterwards, the geometric mean of both reference genes' RQ values was calculated, and based on this, relative genes expression of target genes was calculated, log(2) transformer, averaged over technical duplicates and plotted in GraphPad Prism.

***Life cell imaging in vitro***

SiR-actin and the efflux pump inhibitor Verapamil were purchased as a kit (Spirochrome). Labeling of microglia was performed modifying the manufacturer's recommendations: For SiR-actin labeling, cells were treated with 1 µM SiR-actin and 10 µM verapamil for 3 h at 37°C and 5% $CO_2$ prior to imaging. The medium was kept on the cells during the imaging. Imaging was performed with the Zeiss LSM980 and a 40x oil objective at 37°C, 5% CO2 and a humidified atmosphere.

***Cell morphology in vitro***

Morphology of primary microglia expressing tdTomato was analyzed using the FracLac plugin in Fiji (https://imagej.net/ij/plugins/fraclac/FLHelp/Introduction.htm), determining circularity, density, span ratio and fractal dimensions. Data were plotted in GraphPad Prism and a two-tailed unpaired t-test was performed for statistical analysis.

***Microglia migration in vitro***

Cells expressing tdTomato were recorded for 3 hours at an interval of 1min with an epifluorescent microscope (humified atmosphere, 5% $CO_2$, 37°C).

Videos were analyzed with the „trackMate" plugin for Fiji [1] with the following parameters: 50 µM diameter for thresholding object, 0 quality, DoG detector, LAP tracker. Cells that displayed multiple tracks were excluded or the track that accurately refelects cell movement was chosen for analysis.
